# Root Microbial Functions and Robust Network Drive High-yielding Canola Genotype

**DOI:** 10.1101/2025.04.11.648446

**Authors:** Yunliang Li, Steven D. Mamet, Sally Vail, Steven D. Siciliano

## Abstract

**Highlight:** A high-yielding canola genotype’s productivity is linked to specifically recruited root bacteria, enriched carbon and phosphorus metabolism genes, and a robust microbial network.

Plants and their root microbiomes have co-evolved complex relationships that influence growth and development. While plant genotype is known to shape root microbiomes, a detailed understanding of this interplay remains limited. We used shotgun metagenomic sequencing to examine the composition, function, and microbial association networks of root bacterial communities in two *Brassica napus* (canola) genotypes with contrasting yields: NAM-23 and NAM-30. Root samples were collected at three developmental stages (vegetative, flowering, and maturation) across three field sites. Growth stage significantly influenced alpha diversity and community structure, but not KEGG pathway functions. Genotype had minimal effect on overall diversity and function, but specifically influenced the recruitment of certain bacterial taxa and the topology of microbial association networks. NAM-23, the high-yielding genotype, was enriched in plant growth-promoting bacteria (*Rahnella*) and genes related to carbohydrate and phosphorus metabolism. Additionally, NAM-23 exhibited a more robust and connected microbial network, with higher degree, betweenness, clustering coefficient, and more genotype-specific hub taxa, suggesting enhanced resilience to environmental stress. Our findings provide a high-resolution view of genotype-specific interactions with the root microbiome, highlighting key microbial features associated with high yield. These insights support microbiome-informed strategies for crop improvement in sustainable agriculture.

## 1. Introduction

Plants and their root-associated microbiomes have forged a complex relationship through millions of years of co­evolution that has been pivotal in plant adaptation and diversification. This intricate interplay is central to microbiome engineering, which aims to manipulate microbial communities to improve plant growth, health, and resilience, thereby contributing to more sustainable agricultural practices. Elite genotypes of existing crops provide a unique opportunity to identify key microorganisms and microbial functions inadvertently selected during crop breeding that support plant growth. Here, we focused on how bacterial carbon (C), nitrogen (N), and phosphorus (P) functional pathways differed between two elite genotypes of Canola.

The impact of root-associated microbiomes on host plants is multifaceted and significant. For instance, soybean root inoculation with synthetic communities has been shown to enhance plant growth, nutrient uptake, and yield under varying N and P conditions (Wang et al., 2021). Additionally, artificially assembled microbial communities, along with those naturally recruited by resistant plants, have exhibited considerable disease resistance (Li et al., 2021; Yin et al., 2021). Microbial communities adapted to salinity and drought have been found to improve plant tolerance to these abiotic stresses (Schmitz et al., 2022; Fan et al., 2023).

Plant species, genotype, and plant growth stages influence the assembly and functionality of the root-associated microbiome. Plant species are major determinants in shaping their specific root-associated microbial communities (Ofek et al., 2014; Byers et al., 2023; Ji et al., 2023). Within a species, the effect of genotype on the root-associated microbiome is not consistently significant (Wagner et al., 2016; Barraza et al., 2020; Beschoren da Costa et al., 2022; Li et al., 2023b). The inconsistency might be related to the number and specific genetic loci harbouring mutations across different genotypes, as a particular correlation was identified between plant loci and distinct subsets of the microbiome (Horton et al., 2014; Deng et al., 2021). For instance, *chalcone synthase colorless2* (*C2*), an important gene in flavonoid biosynthesis in maize, is critical for enriching *Massilia*, promoting maize growth and N acquisition (Yu et al., 2021). Similarly, the *NRT1.1B* locus, encoding a nitrate transporter and sensor, contributed to the recruitment of more N-metabolizing bacteria in *indica* rice varieties than *nrtl.lb* mutant-containing *japonica* varieties, which partially accounts for the differences in N use efficiency between the two genotypes (Zhang et al., 2019). Moreover, the dynamics of the root-associated microbiome are significantly influenced by morphological and physiological changes during plant growth, such as alterations in root architecture (Herms et al., 2022), root exudation (Chaparro et al., 2014), and nutrient demand (Pantigoso et al., 2022). Environmental factors, including soil quality (Chen et al., 2019), moisture levels (Fan et al., 2023), temperature (Campisano et al., 2017), and biotic stress (Carrion et al., 2019), are also key drivers of the composition, structure, and function of the root-associated microbial community.

Crop breeding has created elite genotypes that exhibit superior performance across various agronomic traits, encompassing yield productivity and resistance to abiotic and biotic stresses (Reeves and Cassaday, 2002). The variation of genetic background within these elite genotypes influences their metabolisms. Consequently, the varied metabolite profile could impact the selection of microorganisms from the soil (Pang et al., 2021). These genotype-preferred microbes could serve as beneficial partners, playing a pivotal role in enhancing the overall performance of these elite genotypes. While root-associated microbiome profiles among different crop genotypes have frequently been documented (Simonin et al., 2020; Favela et al., 2021; Li et al., 2023b), there is a notable scarcity of reports regarding microbial variations at the functional level. This gap in research is particularly evident when considering distinctions specific to various growth stages.

Microbial composition, structure, and function analyses can offer crucial fundamental information. However, the impact of the microbial community on the host may largely depend on their associations. It is akin to using the same Lego blocks to construct various Lego figures, the outcome of which hinges on the manner of their connections. Microbial co-occurrence networks can provide this extra-dimensional information, which enables us to understand microbial structure better (Fuhrman, 2009; Faust and Raes, 2012). Network centralities—such as degree, closeness, betweenness, and eigenvector—along with hub taxa, describe the properties of microbial co-occurrence networks, reflecting the interactions between the host and microbial communities (Agler et al., 2016; Duran et al., 2018; Hevey, 2018). However, more documentation is needed on the impact of plant genotype on the root microbial co-occurrence network.

In the present study, we selected two canola genotypes, i.e., NAM-23 and NAM-30, from a nested association mapping (NAM) population parental panel which were evaluated in Canada in recent years (Taye et al., 2020; Mamet et al., 2021; Morales Moreira et al., 2021; Singh et al., 2021; Williams et al., 2021; Zhang et al., 2021; Ebersbach et al., 2022; Li et al., 2023 a, b). These two canola genotypes were selected because they significantly differ in yield performance (Williams, 2023). We set up the field trial at three experimental sites and collected roots associated with soil at three growth stages, i.e., vegetative, flowering, and maturation stages. We explored the composition and function of root microbial communities, as well as microbial co-occurrence network. Our hypotheses are as follows: (1) Differences in the yield productivity of canola genotypes are reflected in the relative abundance of microbial taxa, enzymes, clusters of orthologous groups (COGs), and Kyoto encyclopedia of genes and genomes (KEGG) pathways. (2) Microbial metabolisms of carbon (C), nitrogen (N), and phosphorus (P) could drive yield variations between canola genotypes. (3) Genotype-specific root microbial network contributes to canola yield performance.

## 2. Materials and Methods

### 2.1. Site description

The experimental site details have been comprehensively outlined in our previous paper (Li et al., 2023a). Field trials were conducted at the Agriculture and Agri-Food Canada (AAFC) Llewellyn Research Farm (52.1718° N, 106.5052° W) in Saskatchewan, Canada, in 2016 and 2017. Additionally, trials were carried out at two other AAFC Research Farms, Scott (52.3574° N, 108.8400° W) and Melfort (52.8185° N, 104.6027° W), in Saskatchewan in 2017. The three experimental sites have different soil types and properties (Li et al., 2023a).

### 2.2. Experimental design and data collection

NAM-23 and NAM-30 were among 16 canola genotypes selected for exploring the profile of root-associated microbiome using amplicon metagenomic sequencing method (Li et al., 2023a). We picked these two genotypes that showed significant differences in yield productivity for further functional exploration in root microbiomes using the shot-gun metagenomic sequencing method. Across 4 station-years, NAM-23 yielded an average of 2857 kg/ha, which was significantly higher than the average yield of NAM-30 (2038 kg/ha) (Fig. S1). The details of the field trial design were described in a previously published data paper (Bazghaleh et al., 2020). The field trial was a randomized complete block design with three replicates. In 2016, the experiment was set up at Llewellyn Research Farm in Saskatoon, SK, Canada. Roots with attached soil were collected at weeks 3, 6 and 9 starting 18 days after seeding, representing three typical development stages, i.e., vegetative, flowering and maturation stages. Three plants were randomly collected from the inner rows of the plot using a sterile hand trowel to a 10 cm depth. These individual plant samples were composited to form a representative sample of each respective plot. Samples were transported to the laboratory on ice, stored overnight at 4°C and processed the following day. Roots with tightly adhering soil were shaken at 180 rpm for 15 minutes in sterilized 0.05 M NaCl solution. Following the agitation process, the root tissue underwent a sequence of steps: it was flushed with tap water, rinsed with sterilized water, finely chopped using a sterile scalpel, and finally stored at −80°C in preparation for DNA extraction. In 2017, our study was extended to include two additional locations: Scott and Melfort. The same root collection method was applied at the three experimental sites. To calculate the canola yield, plot harvests were carried out using a plot combine, with paired plots located directly adjacent to each microbiome plot that underwent destructive sampling.

### 2.3. DNA extraction, library preparation, and sequencing

DNA was extracted from 50 mg root tissue employing a Qiagen PowerPlant extraction kit (Hilden, Germany) following manufacturer instructions. The extracted DNA samples were then sent to the Next Generation Sequencing Facility at the University of Saskatchewan and stored at - 80°C until library preparation. Before initiating the library preparation process, DNA concentrations were quantified using Qubit dsDNA High Sensitivity assay (Thermofisher), and quality was assessed using Tapestation 4150 genomic DNA ScreenTape assay (Agilent). Libraries were prepared using an Illumina DNA preparation kit (CAT: 20060060). Before segmentation, a total of 10 ng DNA input was normalized to 30 ml using nuclease-free water. Library quantification was performed using the Qubit assay, and the quality was assessed using the Tapestation 4150 High Sensitivity D1000 ScreenTape system. Twenty-four barcoded libraries were pooled at equimolar concentrations (4 nM) and spiked with 1% PhiX control. Finally, libraries were sequenced using a Nextseq 550 High Output v2.5 (300 cycles) to generate 149 bp paired-end reads.

### 2.4. Bioinformatics

#### 2.4.1 Metagenome reads trimming, decontamination, and taxonomic assignment

The quality and integrity of raw sequencing read were checked using FastQC (v0.11.9) (Lo and Chain, 2014). Adapters in raw reads were trimmed by Trimmomatic (v0.39) (Bolger et al., 2014) and host genomes (GCF_020379485.1, GCF_000686985.2, GCA_026770255.1, GCA_026770265.1) and the spiked Phix genome (GCF_000819615.1) were removed using Bowtie2 (v2.5.0) (Langmead and Salzberg, 2012) wrapped in the Kneaddata (v0.12.0) (https://github.com/biobakery/biobakery). Taxonomic assignment of the clean reads was performed using Kraken2 based on a microbial database built from bacterial, archaeal, fungal, viral, and protozoan Refseq in Mar. 2023 (Wood and Salzberg, 2014; Wood et al., 2019). Bracken was then applied to estimate the abundance of microbial taxa (Lu et al., 2017).

#### 2.4.2 Metagenome assembly and binning

For each sample, quality-controlled reads were individually assembled using metaSPAdes (Nurk et al., 2017), and the taxonomic assignment and abundance estimation of the assembled contigs were performed using Kraken2 and Bracken. The metagenomic assembled contigs were binned using MetaBAT2 (v2.15), with the minimum length of the contig set at 2000 bp. The metagenome-assembled genomes (MAG) were assessed and taxonomically classified using GTDB-TK-based MDMcleaner (v0.8.7) (Vollmers et al., 2022).

#### 2.4.3. Functional annotation and quantification

Gene prediction and protein annotation for metagenomic assemblies were performed using Prokka (v1.14.6) (Seemann, 2014), which utilizes the Prodigal (v2.6.3) program (Hyatt et al., 2010). To quantify the predicted genes, clean reads were mapped back to the metagenomic assemblies using bowtie2 (v2.5.0) to generate .sam files. These .sam files were used for constructing highly efficiency indexed files using SAMtools (v1.6) (Danecek et al., 2021). These indexed files were used to calculate the coverage of predicted genes using prokkagff2bed.sh, BEDTtools (v2.31.0) (Quinlan and Hall, 2010), and get_coverage_for_genes.py after the removal of sequencing read duplicates utilizing MarkDuplicates from the Picard Toolkit (https://broadinstitute.github.io/picard/). The scripts, prokkagff2bed.sh and get_coverage_for_genes.py, were developed by the Environmental Genomics group at SciLifeLab Stockholm. Annotated genes and their abundances were extracted based on Enzyme Commission (EC) numbers from .gff files generated by Prokka. The ratios of annotated genes in each sample were between 13.3% and 30.3% (Fig. S2). The abundance of a specific KEGG pathway was calculated based on the abundance of EC numbers linked to this KEGG pathway using KEGGREST (v1.40.1) (Tenenbaum, 2023). The annotation and relative abundance of genes involved in C, N and P cycling metabolism were analyzed using the CAZyme (Zheng et al., 2023), NCycDB (Tu et al., 2019) and PCycDB (Zeng et al., 2022) databases, respectively. These databases are comprehensive and accurate resources for studying genes related to C, N and P metabolism. The details of each analysis process are as follows. The required data files were downloaded and configured using the CAZyme annotation protocol. Subsequently, PROKKA-predicted protein sequences underwent annotation using the HMMER vs dbCAN HMMdb method in the run_dbcan tool. Trimmed reads were then mapped to gene coding sequences to estimate the abundance of CAZyme families. Finally, the abundance was normalized using dbcan_utils with the Transcripts Per Million (TPM) method (Zheng et al., 2024). Following the guidelines and the tool *NCycProfiler.PL* provided by NCycDB, the merged reads were aligned to the database, NCyc_100_2019Jul.faa, using diamond v2.0.15 (Buchfink et al., 2021). Subsequently, the relative abundance of each gene was normalized to the minimum count of merged reads across all samples via random subsampling. According to the instruction of PCycDB, the assembled contigs of each sample were aligned to the database, PCycDBvI.l.faa, using diamond v2.0.15, followed by the extraction of P-cycling genes using the provided script *filter_Generate_ORF2gene.py* with a stricter cutoff (-s 70 - cov 25). The relative abundance of P-cycling genes was further calculated based on the relative abundance of contigs generated by Salmon v0.13.1(Patro et al., 2017) using the provided script *Coverage_get.py*. In addition, we extracted PROKKA-predicted coding DNA sequences (CDS) related to C or P metabolism from each sample by blasting against the CAZyme and PCycDB databases. We then integrated the CDS abundance and taxonomy to calculate the genus-level abundance of C and P metabolism-related CDS in each sample.

### 2.5. Statistical analyses

Bacterial abundance at different taxonomic levels was extracted from the Bracken output file. In the relative abundance analysis of bacterial community, the top 9 bacterial phyla were selected, while the rest were summed as others. Alpha diversity (Shannon index) was calculated using alpha_diversity.py in KrakenTools after cumulative sum scaling (CSS) transformation of species abundance data (Lu et al., 2022). Analysis of variance (ANOVA) was applied to determine the significant effect of the canola genotype and growth stage on the Shannon index based on a linear mixed-effect model. Redundancy analysis (RDA) was used to test the influence of environmental factors (site and year), canola genotype, and growth stage on bacterial community structure and function. The bacterial community at genus level was used for RDA after filtrations of rare genera (prevalence < 0.5), and RDA at the levels of enzyme and KEGG pathway annotated from the bacterial community was applied after filtration of rare enzymes and KEGG pathways (prevalence < 0.25). Hellinger transformation was applied to the three sets of bacterial data for RDA. Package pheatmap (v1.0.12) (Kolde, 2019) was applied to describe the relative abundance of KEGG pathways in the samples at specific conditions, i.e., site, year, genotype, and growth stage.

To detect the differential abundance in bacterial genera, enzymes, and COGs between NAM-23 and NAM-30, we applied three algorithms: LinDA (Zhou et al., 2022), ancomBC2 (Lin and Peddada, 2020a, 2024), and LEfSe (Segata et al., 2011), and integrated their outcomes. As LinDA and ancomBC2 have been reported to be more stringent than LEfSe (Lin and Peddada, 2020b; Zhou et al., 2022), an alpha value of 0.1 was set for LinDA and ancomBC2, whereas an alpha value of 0.05 and an lda threshold of 2.5 were set for LEfSe. For all three methods, feature prevalence was set to 0.1. We selected the plant­growth-promoting rhizobacteria (PGPR) and plant pathogens included in the differentially abundant bacterial genera to further explore which bacterial species or strains might affect canola yield productivity. Subsequently, a Wilcoxon test was applied to detect differences in their relative abundance between NAM-23 and NAM-30. To test the differential abundance of C, N and P metabolism-related genes between NAM-23 and NAM-30, only those genes that occurred in more than half of the samples were considered. Specifically, 54 out of 346 genes associated with C metabolism, 47 out of 66 genes involved in N metabolism genes, and 120 out of 131 genes related to P metabolism were selected for differential abundance analysis using the Wilcoxon test.

The microbial co-occurrence and differential networks were constructed and analyzed using NetCoMi (v1.1.0). Bacterial genera were filtered using two criteria: (1) Each genus frequency is higher than 0.8; (2) At least in one sample, the genus abundance is higher than 0.05% of the sample’s total count. There, 226 out of 1510 genera were retained, and then the filtered bacterial genera table was split into two tables by canola genotype, i.e., NAM-23 and NAM-30. Data sparsity was not an issue in our concerns because only a few zero values (≤ 10) were presented in each bacterial genus table after filtration. Zero values were replaced using the compositionally-aware method of multiplicative replacement. Pearson correlation was applied to calculate the association matrix after bacterial genus data’s centred log-ratio (CLR) transformation. Correlation coefficients with magnitude > 0.7 or < −0.7 were selected. To better explain the properties of the microbial co-occurrence network, the related terminologies were described as follows: (a) Degree centrality is a measure that quantifies the importance of a node (bacterial genus) in a network based on the number of connections it has; (b) Closeness centrality is a measure that quantifies how close a node is to all other nodes in a network; (c) Betweenness centrality is a measure that quantifies the importance of a node in controlling the flow of information or resources between other nodes in a network; (d) Eigenvector centrality measures the influence of a node in a network, considering both the node’s direct connections and the connections of its neighbours; (e) Clustering coefficient is a network metric that measures the extent to which nodes in a network tend to cluster together (Hevey, 2018; Peschel et al., 2021). Two distinct selection criteria were applied to identify the hub taxa: one was that the genera selected must be within the top 15% (quantile 85) of normalized centralities of degree, betweenness, and closeness simultaneously (Agler et al., 2016) and the other was within the top 5% (quantile 95) of normalized eigenvector centrality (Csardi and Nepusz, 2005).

Centralities of degree, betweenness and closeness depicted node properties, and the node with high centrality indicates its prominent position in the network (Agler et al., 2016; Peschel et al., 2021). A node with high eigenvector centrality is connected to other nodes that are themselves well-connected, indicating a central and influential position in the network (Ruhnau, 2000; Peschel et al., 2021). The “Cluster_fast_greedy” algorithm was applied to identify potential functional groups. Differential taxa associations between NAM-23 and NAM-30 were determined using Fisher’s z-test, and *p-values* were adjusted using the Local False Discovery Rate (lFDR) (Efron, 2005).

## 3. Results

### 3.1. Composition, structure and function of root bacterial community responding to various factors

The relative abundance patterns of the root bacterial community varied across sites, years, canola genotype and growth stage (Fig. 1a and b). Pseudomonadota and Actinomycetota are the dominant bacterial phyla, accounting for over 80% of bacteria under all conditions. The root microbial alpha diversity (Shannon Index) of NAM-30 tended to be higher than that of NAM-23. Their difference is close to the significant level (*p* = 0.066) (Fig. 1c). Growth stage had a considerable effect on root microbial alpha diversity, which was indicated by the higher root microbial Shannon index at week 9 than at week 3 (Fig. 1d). RDA of the microbial community suggested that site, sampling year and growth stage are the significant factors in shaping bacterial community at the genus level (Fig. 1e). In contrast, sampling year is the main driver to the variation of microbial gene function (Fig. 1f). However, none of such selected factors had a significant influence on the higher hierarchical function unit, KEGG pathway.

**Figure 1.**
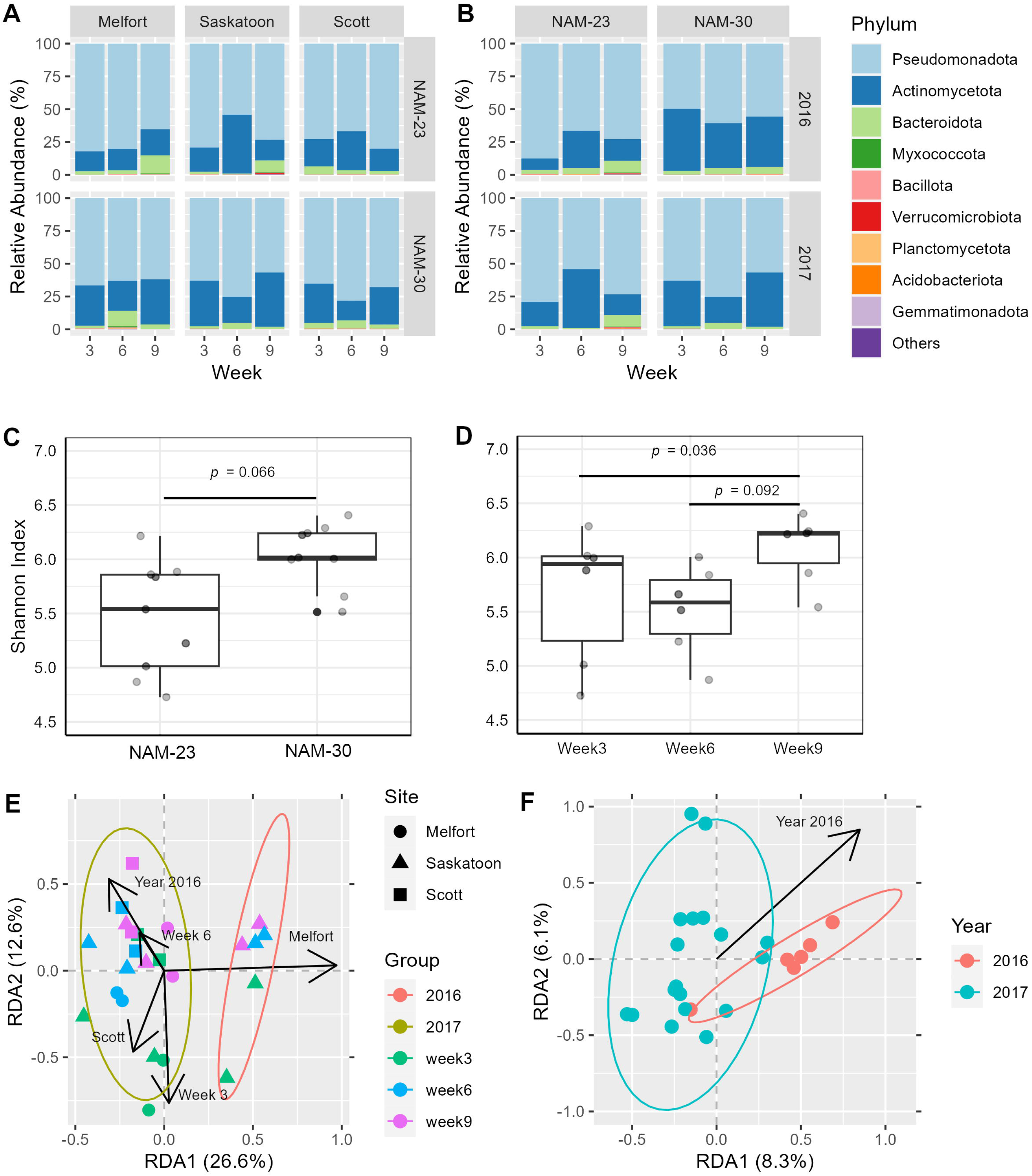
Profile and diversity of root bacterial community across experimental sites, years, canola genotypes and growth stages. **a**, Relative abundance of root bacterial community at phylum level in NAM-23 and NAM-30 at three growth stages (week3, 6, and 9) across three sites, i.e., Melfort, Saskatoon, and Scott across in 2017, and **b**, between 2016 and 2017 in Saskatoon; **c**, Shannon index of bacterial community between NAM-23 and NAM-30 and **d**, at three growth stages, their significant differences were determined by ANOVA; **e**, Redundancy analysis (RDA) of factors, i.e., site, year, genotype and growth stage, and root bacterial community; **f**. RDA of microbial enzymes.

### 3.2. Differentially abundant root bacteria, their proteins, and metabolic pathways between NAM-23 and NAM-30

To partially mitigate the large p, small n problem inherent in metagenomic analysis, we used three different algorithms to assess relative abundance. Specifically, we selected three methods—LinDA, ancomBC2, and LEfSe—for identifying differentially abundant ECs, KEGG pathways, COGs, and bacteria between NAM-23 and NAM-30. LEfSe identified 16 differentially abundant bacteria at the genus level, while LinDA and ancomBC2 did not detect any significant bacteria (Fig. 2a and d). Interestingly, *Rahnella* bacteria were highly enriched in NAM-23. In contrast, the remaining 15 bacterial genera were more abundant in NAM-30 (Fig. 2a). Fifteen out of 192 species or strains included in these differentially abundant bacterial genera were reported as plant-growth­promoting rhizobacteria (PGPR) or plant pathogens (Table S1), and 8 out of the 15 species or strains were different in relative abundance between NAM-23 and NAM-30.

**Figure 2.**
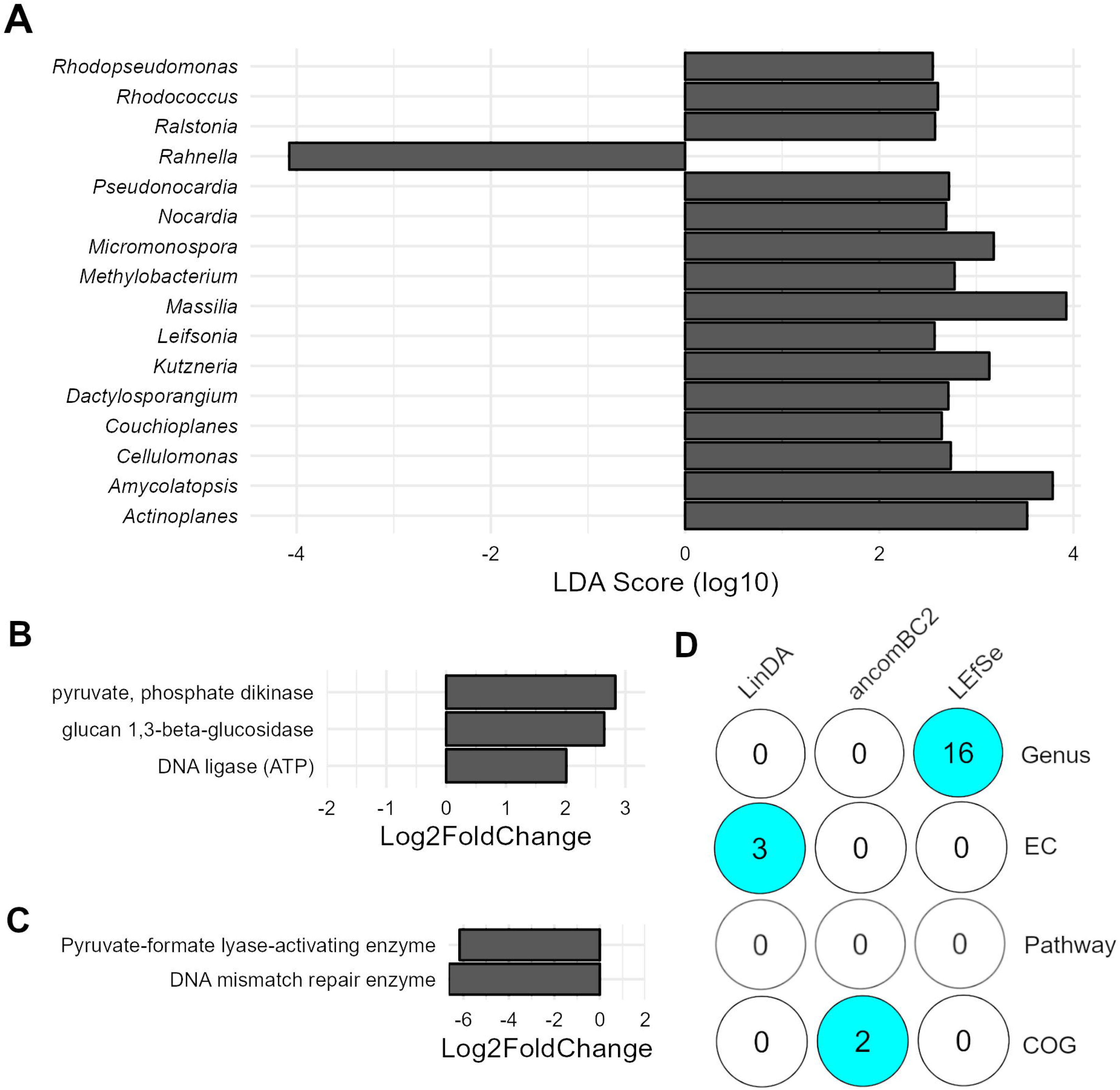
Differential abundance analysis **a.** bacterial genera NAM-23 vs. NAM-30 using LEfSe (lda = 2.5); **b.** microbial enzymes NAM-23 vs. NAM-30 using LinDA (α = 0.1); **c.** Cluster of Orthologous Groups (COG) NAM-23 vs. NAM-30 using ancomBC2 (α = 0.05)); **d.** Numerical methods used and number of identified differentially abundant features: Bacterial Genus, Enzyme Commision (EC), KEGG Pathway (Pathway) and Cluster of Orthologous Groups (COG).

Specifically, 3 PGPR species in *Rahnella* were enriched in NAM-23, while 2 PGPR species in *Micromonospora*, 1 PGPR species in *Leifsonia*, and 2 other plant pathogens were enriched in NAM-30 (Fig. S3). Furthermore, three ECs, i.e., pyruvate phosphate dikinase, glucan 1,3-beta-glucosidase, and DNA ligase identified by LinDA displayed higher abundance in NAM-30 than NAM-23 (Fig. 2b and d). Additionally, two COGs, pyruvate-formate lyase-activating enzyme and DNA mismatch repair enzyme, identified by ancomBC2 were enriched in NAM-23 (Fig. 2c and d). No differentially abundant KEGG pathways were identified.

Between each pair of the three growth stages in both canola genotypes, LEfSe identified more bacterial genera than LinDA and ancomBC2. However, NAM-23 and NAM-30 only had one differentially abundant bacterial genus in common for week 3 vs 6 and week 3 vs 9, respectively (Supplementary Excel). The KEGG pathways of the root microbial community were split into two main clusters based on their relative abundances in each sample. The relative abundances of KEGG pathways in cluster 1 were stable across each sample, while they had large variations in cluster 2 (Fig. S4).

### 3.3. Differentially abundant genes associated with C, N, and P metabolism between two canola genotypes

To further investigate the influence of the canola genotype on root microbial metabolisms, we conducted comprehensive annotations of all metagenomic genes related to C, N, and P metabolisms using specialized databases. We then computed their normalized abundances and performed differential abundance analysis.

Heatmap analyses revealed that a significant portion of microbial genes related to C and P metabolism were more abundant in NAM-23 compared to NAM-30, whereas genes associated with N metabolism showed no such trend (Fig. 3a, c and e). In addition, microbial C (8 out of 54), N (5 out of 47) and P (9 out of 120) metabolism-related genes had statistically significant differences in abundance between NAM-23 and NAM-30. Among these differentially abundant genes, most C (7 out of 8) and P (6 out of 9) metabolism-related genes were elevated in high-yield NAM-23 compared to NAM-30 (P < 0.05), while NAM-30 enriched more N (4 out of 5) metabolism-related genes than NAM-23. The significantly-enriched C metabolism-related enzymes in NAM-23 belong to the glycosyl transferases, glycoside hydrolases, carbohydrate esterase, and auxiliary activities (Fig. 3a and b). Notably, NAM-23 exhibited a higher abundance of *gdh_K00262*, a hub enzyme bridges N and C metabolism by catalyzing the reversible conversion between glutamate and α-ketoglutarate with ammonia assimilation or release, whereas NAM-30 showed elevated levels of *nirA* and *NR*, two essential genes in assimilatory nitrate reduction (Fig. 3c and d). In NAM-23, the P genes of *pyk*, *pstS*, *ppa*, *phoU*, and *gmk*, were elevated (Fig. 3e and f). In addition, we further analyzed the distribution of C and P metabolism-related gene at genus level. The distribution pattern of C metabolism-related genes in NAM-23 was distinct from that in NAM-30. Apart from *Streptomyces*, which showed a comparable abundance to NAM-30, *Stenotrophomonas*, *Serratia*, and *Pseudomonas* were the dominant genera in NAM-23, collectively harbouring over half of the C metabolism-related genes (Fig. S5a). However, the distribution patterns of P metabolism-related genes were similar between NAM-23 and NAM-30. The dominant genera harbouring P metabolism-related genes were the same in both NAM-23 and NAM-30, although a higher proportion of gene abundance originated from these dominant genera in NAM-23 (Fig. S5b).

**Figure 3.**
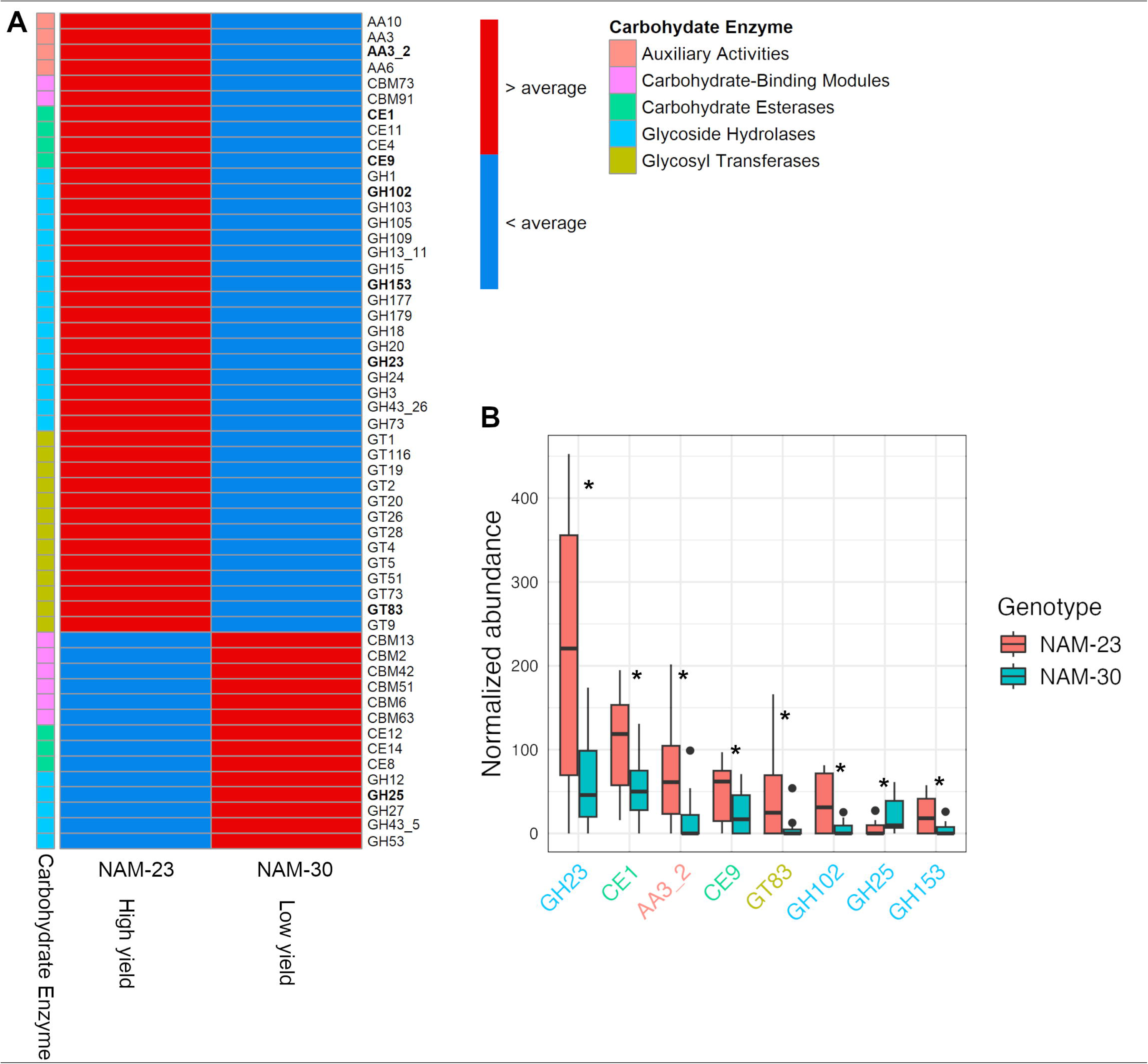
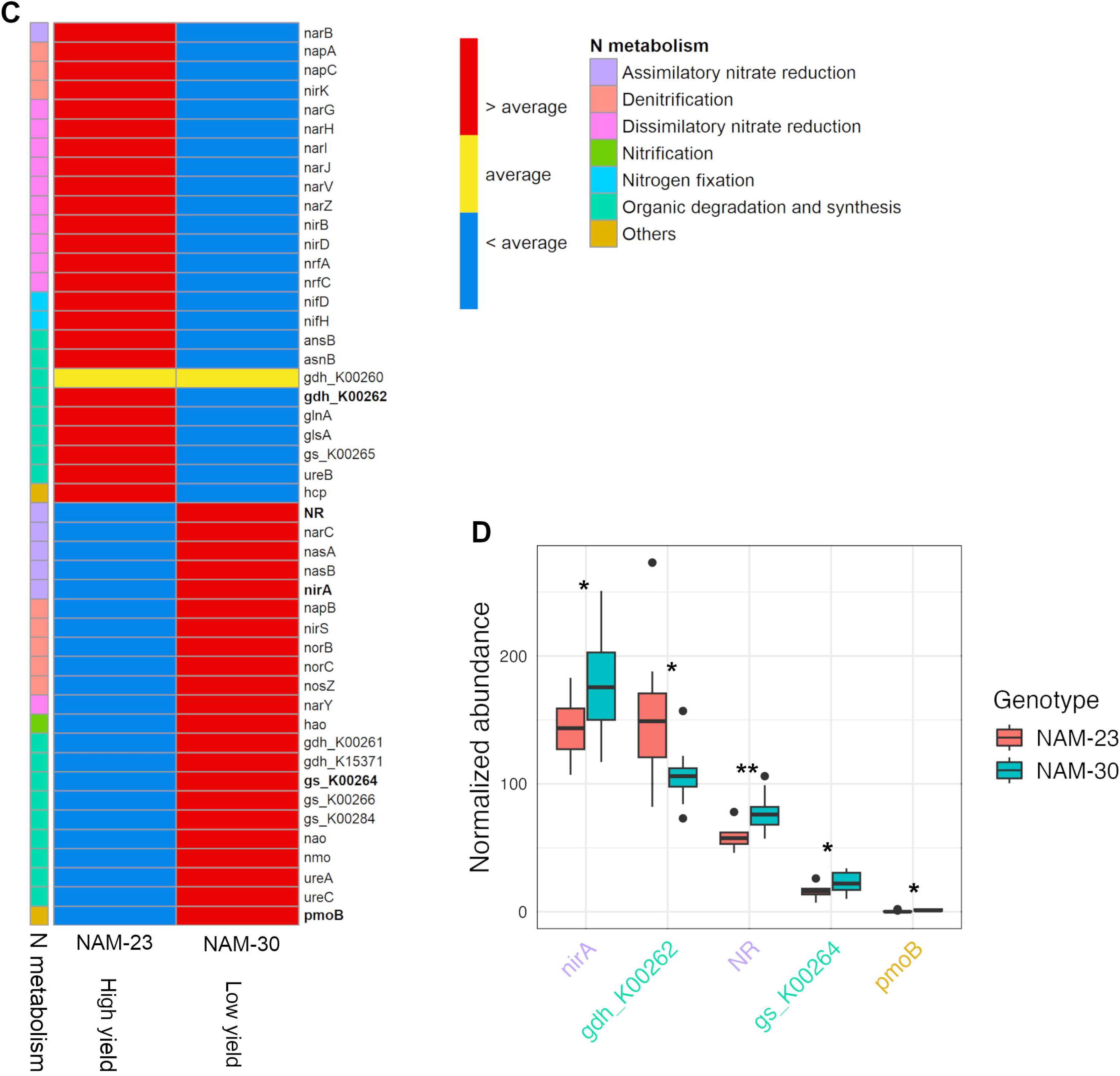
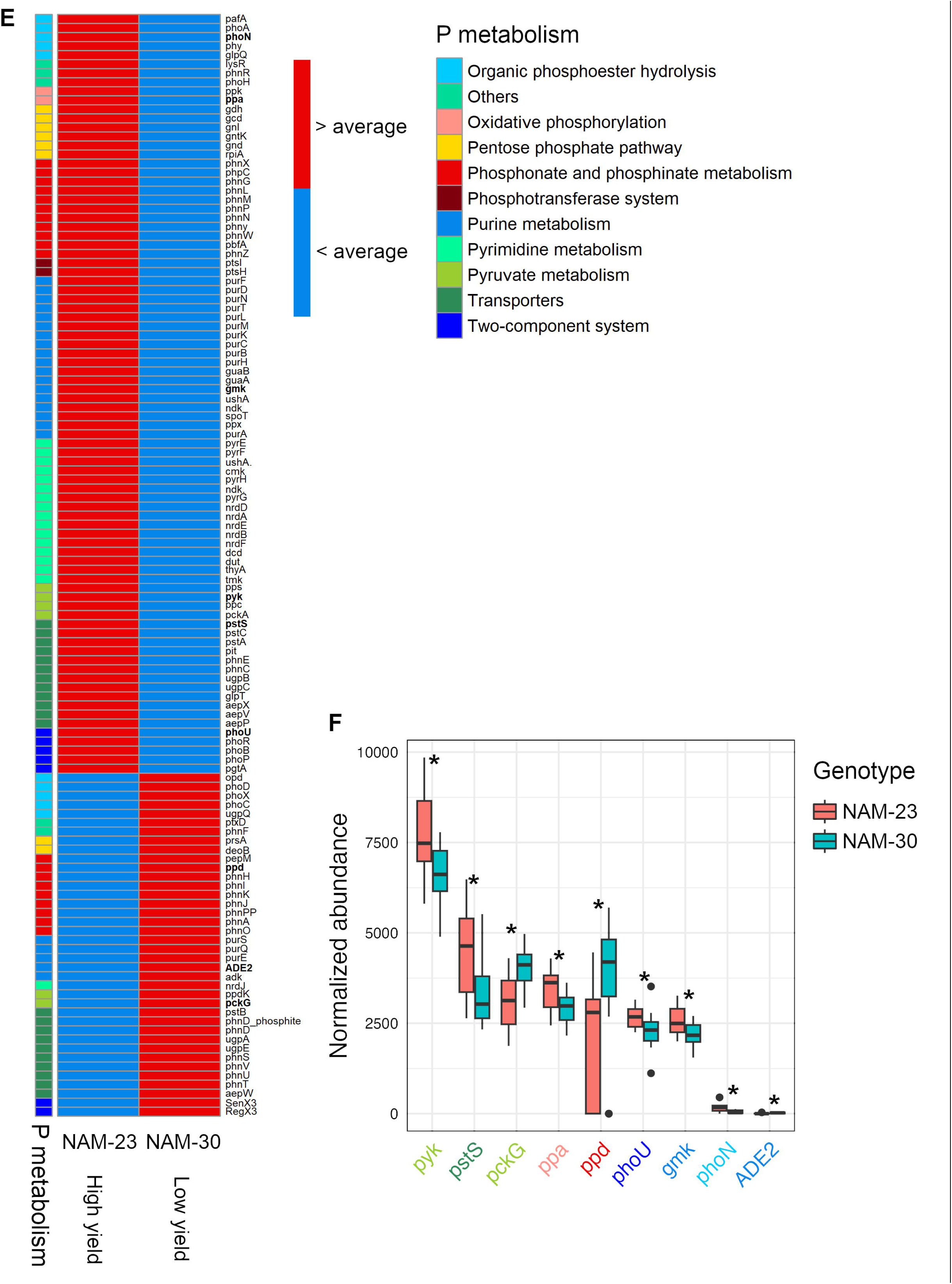
Heatmap and bar-plot showing gene relative abundance between NAM-23 and NAM-30. a, b, carbohydrate metabolism related genes; c, d, nitrogen metabolism related genes; e, f, phosphorus metabolism related genes. Significance was tested using Wilcoxon-test, * indicates *p* < 0.05; ** indicates *p* < 0.01, *p* values underwent no adjustment.

### 3.4. Microbial co-occurrence networks

Although only a few differentially abundant bacterial species were identified between NAM-23 and NAM-30, the topological characteristics of the microbial co-occurrence network differed significantly between the two canola genotypes in terms of degree, betweenness, closeness, and eigenvector centralities, as well as hub taxa and clustering coefficients. Among the top 10 bacterial genera ranked by degree, betweenness, closeness, or eigenvector centrality, there was no overlap in degree or eigenvector centrality between NAM-23 and NAM-30. However, one genus (*Acidovorax*) was shared in betweenness centrality, and two genera (*Melaminivora* and *Ottowia*) were shared in closeness centrality. Additionally, the top 10 degree and betweenness centralities were higher in NAM-23 compared to NAM-30, while closeness centrality was lower in NAM-23 (Table 1).

**Table 1.**
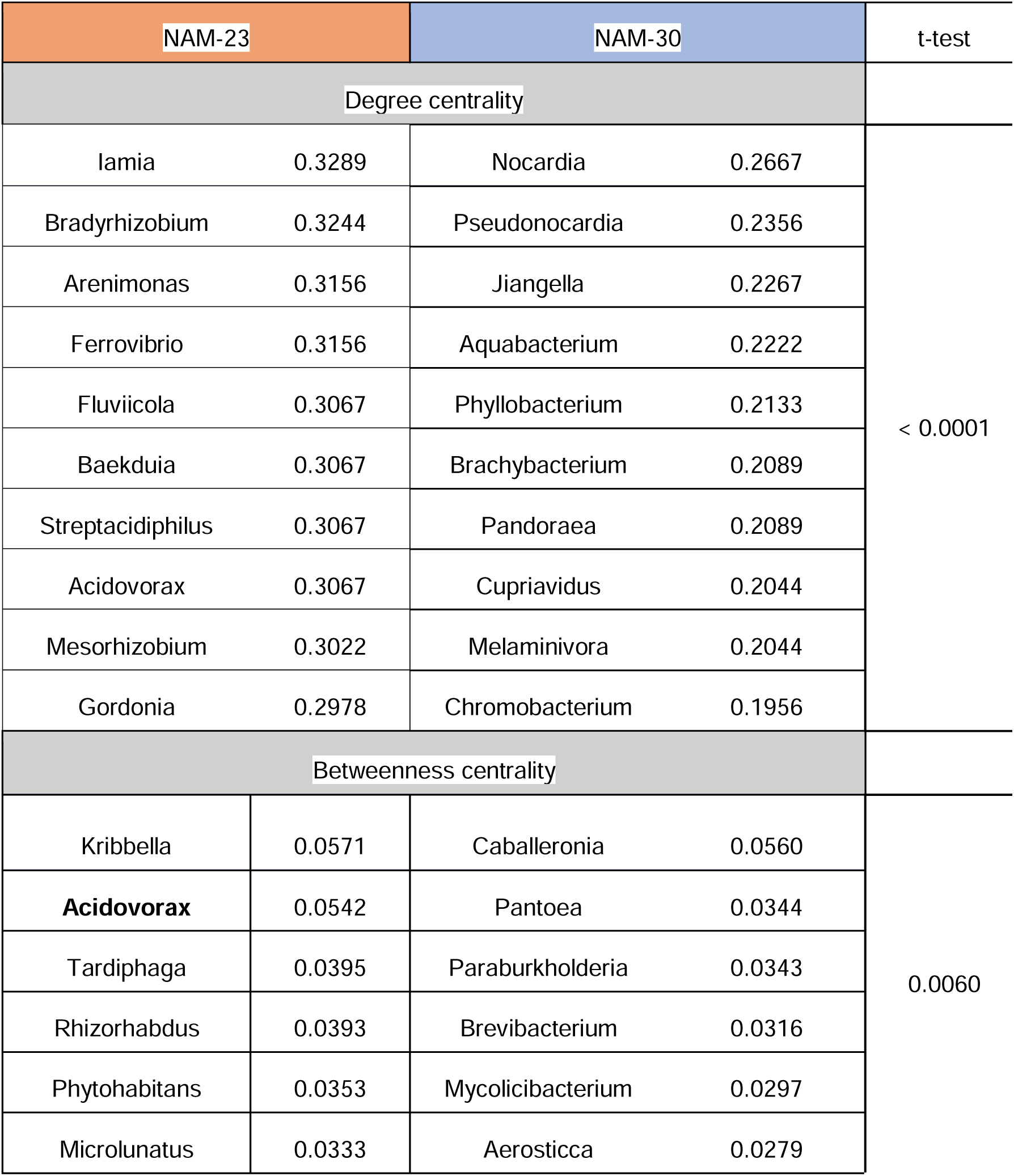
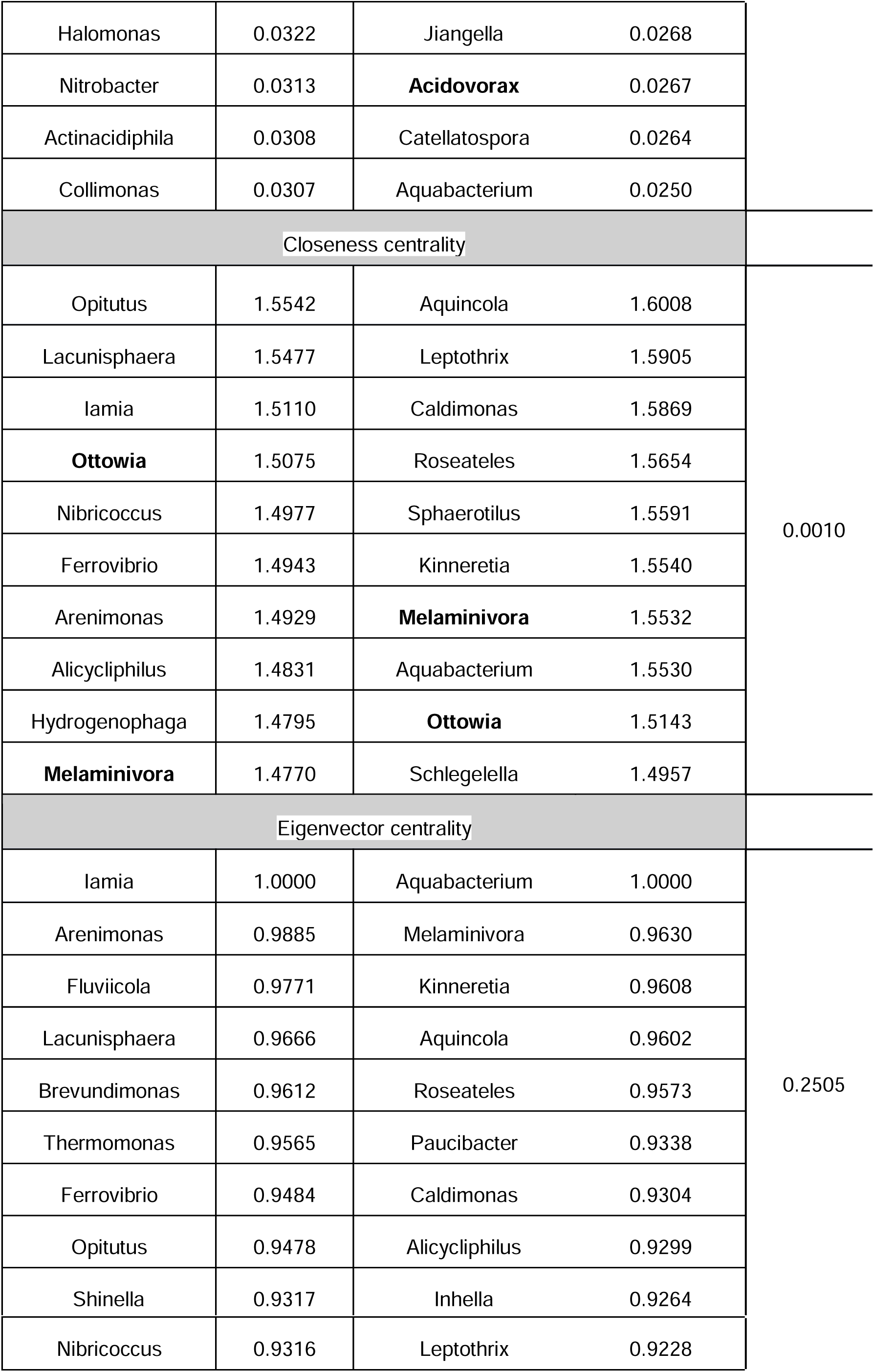
Top 10 genera in each group regarding normalized centralities of degree, betweenness, closeness, and eigenvector in NAM-23 and NAM-30. The centrality difference between NAM-23 and NAM-30 in each group was indicated using t-test. Genera in bold occurred in both NAM-23 and NAM-30.

| NAM-23 |  | NAM-30 |  | t-test |
| --- | --- | --- | --- | --- |
| Degree centrality |  |  |  |  |
| Iamia | 0.3289 | Nocardia | 0.2667 | < 0.0001 |
| Bradyrhizobium | 0.3244 | Pseudonocardia | 0.2356 |  |
| Arenimonas | 0.3156 | Jiangella | 0.2267 |  |
| Ferrovibrio | 0.3156 | Aquabacterium | 0.2222 |  |
| Fluviicola | 0.3067 | Phyllobacterium | 0.2133 |  |
| Baekduia | 0.3067 | Brachybacterium | 0.2089 |  |
| Streptacidiphilus | 0.3067 | Pandoraea | 0.2089 |  |
| Acidovorax | 0.3067 | Cupriavidus | 0.2044 |  |
| Mesorhizobium | 0.3022 | Melaminivora | 0.2044 |  |
| Gordonia | 0.2978 | Chromobacterium | 0.1956 |  |
| Betweenness centrality |  |  |  |  |
| Kribbella | 0.0571 | Caballeronia | 0.0560 | 0.0060 |
| Acidovorax | 0.0542 | Pantoea | 0.0344 |  |
| Tardiphaga | 0.0395 | Paraburkholderia | 0.0343 |  |
| Rhizorhabdus | 0.0393 | Brevibacterium | 0.0316 |  |
| Phytohabitans | 0.0353 | Mycolicibacterium | 0.0297 |  |
| Microlunatus | 0.0333 | Aerosticca | 0.0279 |  |
| Halomonas | 0.0322 | Jiangella | 0.0268 |  |
| Nitrobacter | 0.0313 | **Acidovorax** | 0.0267 |  |
| Actinacidiphila | 0.0308 | Catellatospora | 0.0264 |  |
| Collimonas | 0.0307 | Aquabacterium | 0.0250 |  |
| Closeness centrality |  |  |  |  |
| Opitutus | 1.5542 | Aquicola | 1.6008 | 0.0010 |
| Lacunisphaera | 1.5477 | Leptothrix | 1.5905 |  |
| Iamia | 1.5110 | Caldimonas | 1.5869 |  |
| **Ottowia** | 1.5075 | Roseateles | 1.5654 |  |
| Nibricoccus | 1.4977 | Sphaerotilus | 1.5591 |  |
| Ferrovibrio | 1.4943 | Kinneretia | 1.5540 |  |
| Arenimonas | 1.4929 | **Melaminivora** | 1.5532 |  |
| Alicyclophilus | 1.4831 | Aquabacterium | 1.5530 |  |
| Hydrogenophaga | 1.4795 | **Ottowia** | 1.5143 |  |
| **Melaminivora** | 1.4770 | Schlegelella | 1.4957 |  |
| Eigenvector centrality |  |  |  |  |
| Iamia | 1.0000 | Aquabacterium | 1.0000 | 0.2505 |
| Arenimonas | 0.9885 | Melaminivora | 0.9630 |  |
| Fluviicola | 0.9771 | Kinneretia | 0.9608 |  |
| Lacunisphaera | 0.9666 | Aquicola | 0.9602 |  |
| Brevundimonas | 0.9612 | Roseateles | 0.9573 |  |
| Thermomonas | 0.9565 | Paucibacter | 0.9338 |  |
| Ferrovibrio | 0.9484 | Caldimonas | 0.9304 |  |
| Opitutus | 0.9478 | Alicyclophilus | 0.9299 |  |
| Shinella | 0.9317 | Inhella | 0.9264 |  |

|  |  |  |  |
| --- | --- | --- | --- |
| Nibricoccus | 0.9316 | Leptothrix | 0.9228 |

To identify hub taxa, we employed two independent criteria. Using the 85th quantile as the threshold for degree, betweenness, and closeness centralities simultaneously, six hub taxa were identified in NAM-23 and three in NAM-30 (Table 2), with no overlap between the two genotypes. Notably, only one hub taxon (*Bradyrhizobium*) ranked within the top 5% in relative abundance. The relative abundance of the nine hub taxa was comparable between NAM-23 and NAM-30 (Fig. 4a). Applying the 95th quantile of eigenvector centrality as the threshold revealed 12 distinct hub taxa in NAM-23 and NAM-30, respectively (Table 2). Three main clusters were established in NAM-23, while one more main cluster was shaped in NAM-30 (Fig. 4b). However, two main clusters, including taxa nodes marked in blue and purple, were more compact in NAM-23 than in NAM-30 (Fig. 4b), which was also indicated by the higher clustering coefficient in NAM-23 (0.583 in NAM-23, 0.537 in NAM-30).

**Figure 4.**
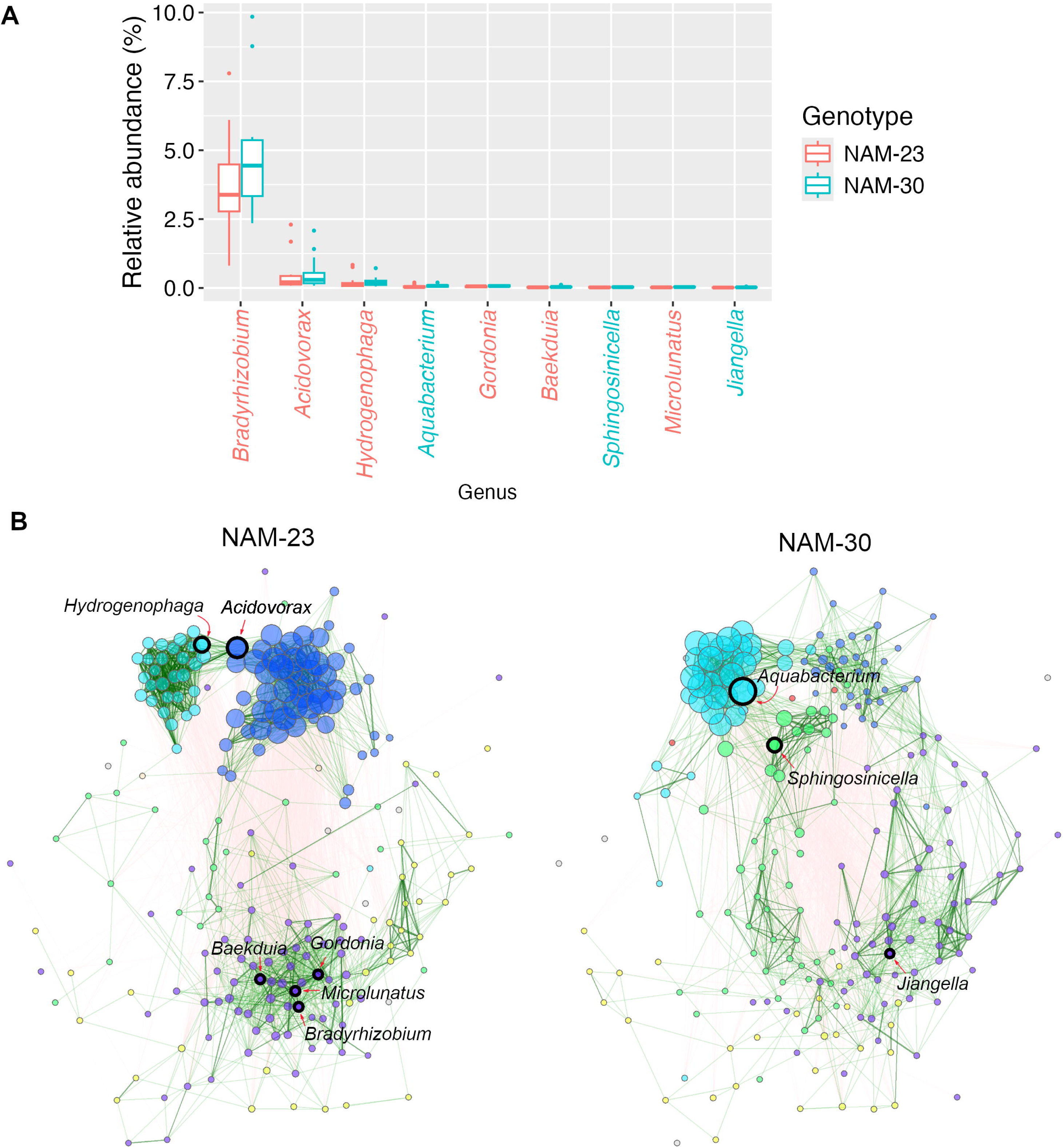
Comparison of hub taxa and bacterial co-occurrence networks between NAM-23 and NAM-30. **a**, Relative abundance of hub taxa in root bacterial community of NAM-23 and NAM-30, genera in different colors on X-axis stand for the hub taxa identified in NAM-23, the rest were identified in NAM-30; number on the top of each bar indicate the rank of relative abundance of the corresponding genus in 226 genera in NAM-23 and NAM-30; **b**, Bacterial co-occurrence network in NAM-23 and NAM-30. Hub taxa were identified by simultaneously considering the top 15% of degree, betweenness, and closeness centralities; hub taxa are highlighted with bold borders; node colors represent clusters; node size is scaled by eigenvector centrality. Green edges correspond to positive estimated associations and red edges to negative ones. The layout of both networks is the same. Nodes that are unconnected in both groups are removed.

**Table 2.**
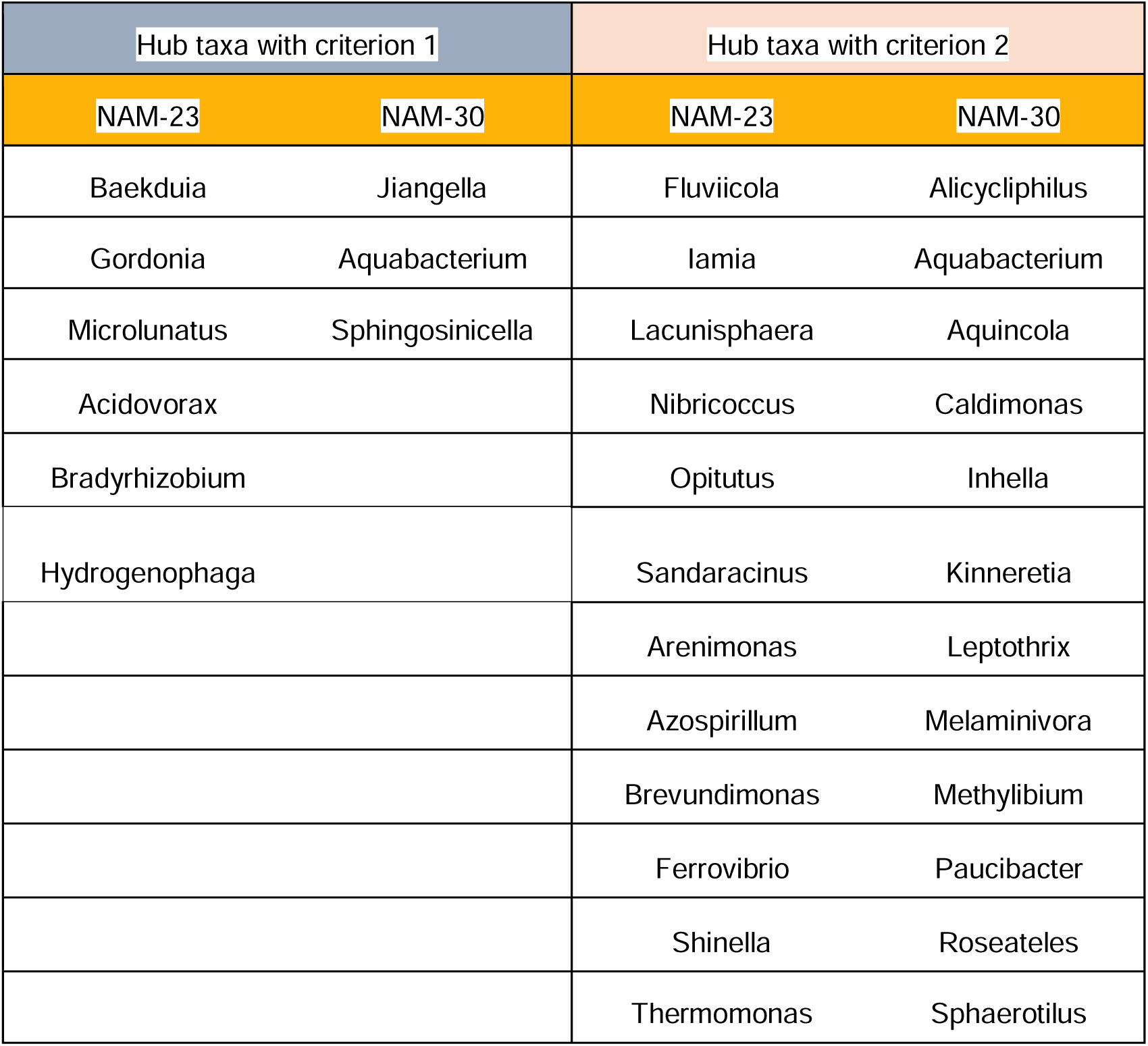
Hub taxa identified in root microbial co-occurrence network of NAM-23 and NAM-30 using two criteria respectively. Criterion 1: genera selected must be within the top 15% (quantile 85) of normalized centralities of degree, betweenness, and closeness simultaneously; Criterion 2: genera selected must be within the top 5% (quantile 95) of normalized eigenvector centrality.

Interestingly, the cluster of taxa nodes marked in purple contained 4 hub taxa in NAM-23 but only 1 hub taxon in NAM-30. However, the hub taxa selected based on eigenvector centrality were exclusively included in the cluster of taxa nodes marked in blue in NAM-23 and the cluster of taxa nodes marked in cyan in NAM-30 (Fig. S6).

To better understand the differing taxa-taxa relationships between NAM-23 and NAM-30, we extracted differential microbial associations. Most differential associations between taxa displayed contrasting patterns between the NAM-23 and NAM-30 networks (Fig. S7). For instance, *Bosea* exhibited a strong positive association with *Rhodopseudomonas* in NAM-23, whereas it showed an opposing trend in NAM-30. *Pandoraea* and *Rhizorhabdus*, *Sphingomonas* and *Massilia*, *Phyllobacterium* and *Saccharothrix* displayed negative associations in NAM-23 but demonstrated positive associations in NAM-30. The networks of NAM-23 and NAM-30 exhibited significantly distinct association structures when considering only differentially associated taxa (Fig. 5). Interestingly, robust associations among *Fluviicola*, *Opitutus*, *Lacunisphaera*, and *Ferrovibrio* disappeared in NAM-30.

**Figure 5.**
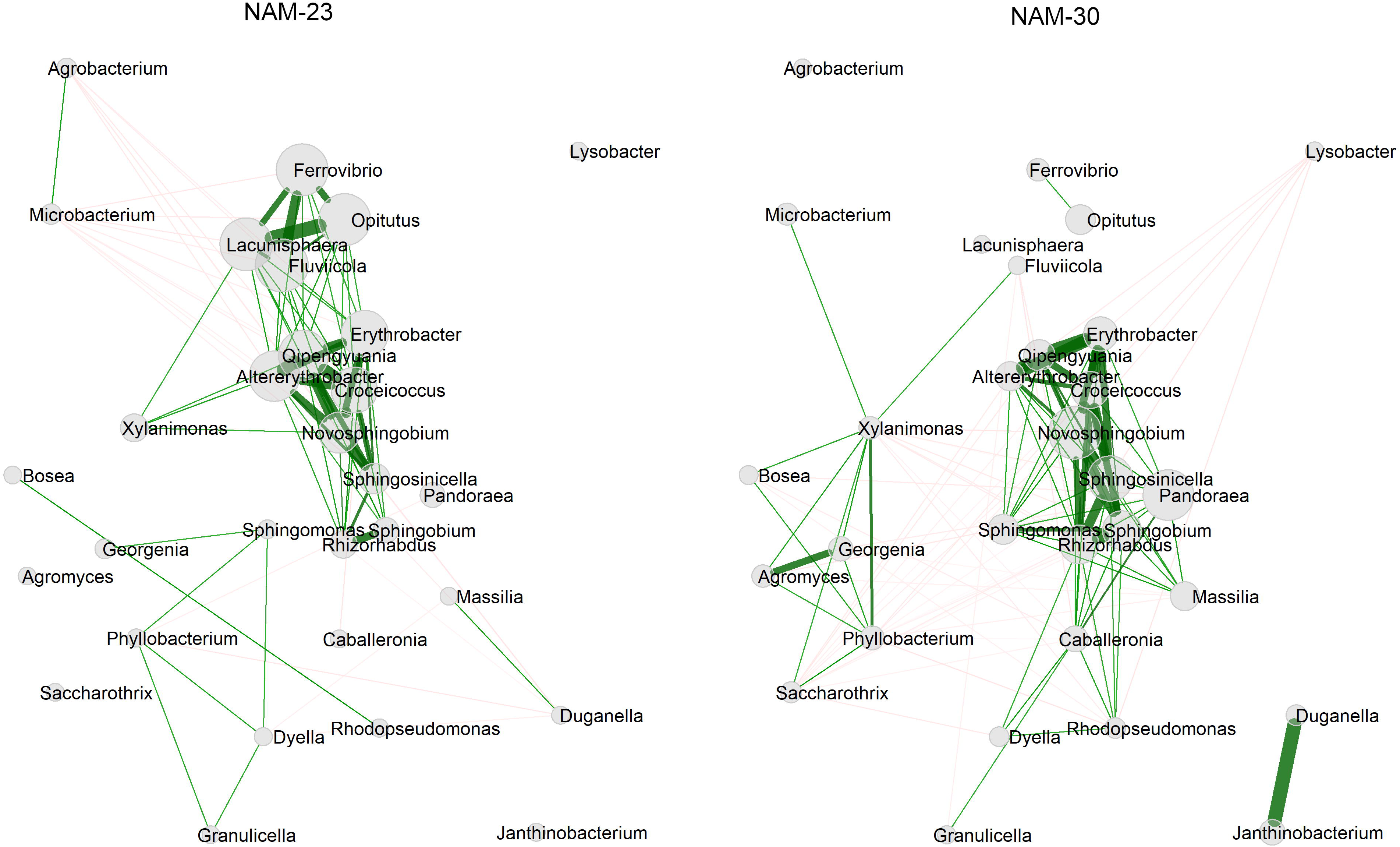
Comparison of bacterial networks constructed using genera with differential association between NAM-23 and NAM-30. The similarities indicating the connection strength of two nodes are used as edge weights. Green edges correspond to positive estimated associations and red edges to negative ones. Node size is scaled by eigenvector centrality. The layout of both networks is the same.

## 4. Discussion

The differences between the high and low yielding canola genotypes was linked to: (i) the enrichment of a specific bacterium, *Rahnella*, and its C and P metabolic pathways as well as (ii) a more complex and stable bacterial network of interactions. We suggest that this combination of a specific beneficial bacterium, coupled with network stability, is one of the rhizosphere traits that contribute to the increased yield of NAM-23.

### 4.1. *Rahnella* bacteria, a potential main contributor to canola productivity

Although we applied three statistical tools to identify differentially abundant microbial taxa, genes, COGs, and KEGG pathways between the two canola genotypes, only few differentially abundant taxa were identified. Our results suggest that the differences between elite canola genotypes have a limited effect on the microbial community at both composition and function levels. There have been reported variations in the influence of plant genotype on root microbial community, ranging from limited to significant effect (Wagner et al., 2016; Barraza et al., 2020; Beschoren da Costa et al., 2022; Li et al., 2023b). However, these effects are less pronounced than environmental factors, such as soil origin (Bonito et al., 2014; Beschoren da Costa et al., 2022). Genome-wide association studies (GWAS) have revealed that most plant genetic loci exert small effects on microbial communities (Bergelson et al., 2021), while certain genomic regions show a robust correlation with microbial abundance and composition (Horton et al., 2014; Deng et al., 2021). This suggests that the extent to which plant genotype influences microbial communities depends on the specific genetic loci The abundance of change in specific bacteria at a fine-scale level between plant genotypes can contribute to their phenotypic variations (Pérez-Jaramillo et al., 2017; Zhang et al., 2019). Among the differentially abundant bacterial genera, NAM-23 enriched *Rahnella*, and most species in *Rahnella* have been widely reported to promote plant growth (Peng et al., 2019; Kong et al., 2022; Xu et al., 2022). Although NAM-30 enriched 3 PGPR, it also recruited more plant pathogens. The synergistic effects of root microbial members in NAM-23 may confer greater benefits to canola yield productivity than NAM-30. Interestingly, our study found that root microbial metagenomes of NAM-23 exhibited enrichment of C and P metabolism genes compared to NAM-30. Previous research has reported that the *Rahnella* genus harbours a rich repertoire of C metabolism and inorganic ion transport genes (Myszka et al., 2023; Wang et al., 2023; ZOUAGUI et al., 2024), and *Rahnella* is widely recognized as phosphate-solubilizing bacteria (Zhang et al., 2018; Landa-Acuna et al., 2023). The consistent trend indicates that the abundant *Rahnella* bacteria strongly contributed to the differences in microbial C and P metabolism-related genes between NAM-23 and NAM-30. Carbohydrate metabolism regulated by bacteria can affect plant growth and defence system by altering soluble sugar level, sugar type and distribution (Su et al., 2024), and *Rahnella*, as phosphate-solubilizing bacteria, can improve the ability of plant P uptake. The superior yield productivity of NAM-23 may partially be attributed to the enrichment of *Rahnella*. In future work, we recommend testing the effect of *Rahnella* inoculant on canola yield productivity in the field.

### 4.2. Understanding canola yield variation through microbial co-occurrence network

Although plant genotype had limited impact on root bacterial community composition and structure, it altered the connectivity and complexity among root bacterial taxa. The root bacterial co-occurrence network of NAM-23 demonstrated elevated network centralities in degree and betweenness, coupled with a higher clustering coefficient than NAM-30. In addition, the main clusters were more compact in NAM-23 than in NAM-30. These findings suggest that NAM-23 harboured a more complex and stable bacterial network. This increased complexity and stability may contribute to enhancing the adaptability of NAM-23 to environmental conditions, ultimately resulting in improved yield productivity across all three experimental sites.

Previous research has shown a correlation between network complexity and crop productivity (Tao et al., 2018; Ji et al., 2022). The heightened network complexity suggests that the relationships between microbial taxa have become more intricate and interconnected. This increased complexity can contribute to functional diversity and redundancy within the microbial community, enhancing the community’s resilience to perturbations. Research has reported that salinity can increase microbial network connections in an anammox system, leading to improved adaptability of the microbial community to elevated salinity (Ya et al., 2021). Compared to conventional farming ecosystems, organic and natural farming ecosystems exhibit higher resilience to abiotic and biotic stresses, potentially attributed to their more complex microbial networks (Banerjee et al., 2019; Tatsumi et al., 2023).

Plant genotype impacted the complexity of the root bacterial network and altered the importance of individual bacterial taxon. Hub taxa selected based on the network centralities of degree, closeness, and betweenness reflected the strength of their connection with other nodes and their ability to connect sub-network. More hub taxa in NAM-23 and no overlapping in hub taxa between NAM-23 and NAM-30 indicate that plant genotype dramatically affected the interaction between bacterial taxa. The abundance of hub taxa was comparable between NAM-23 and NAM-30, indicating that the interaction way of microbial taxa other than their relative abundance determines their role in a network and the function of the microbial community (Tapio et al., 2017; Meng et al., 2022). It was interesting to find that the effect directions of some microbial interactions were opposite between canola genotypes, as shown in the differential network. Microbial interaction is dynamic and depends on context (Coyte and Rakoff-Nahoum, 2019).

Microbes supporting each other based on their metabolites become mutualisms in a specific context. However, the positive relationship might break down and become competitive under limited resource conditions (Hoek et al., 2016). The variation of plant genotype in genetic background likely results in changes in the profile of metabolites, potentially affecting the microbe-microbe interaction.

## 5. Conclusion

Although plant genotype has a limited effect on root microbial composition, structure and function at the community level, the significant changes in the relative abundance of specific microbes and C and P metabolism related genes between plant genotypes potentially contribute to their phenotypic variations. *Rahnella* bacteria have the potential to be considered biofertilizers for canola. Moreover, the features of microbial co-occurrence networks, including network centralities, clustering, and hub taxa, provided an additional dimension of information about microbial associations at the systematic level, which aids in enhancing our comprehension of the intricate relationship between plant genotype and the root microbial community.

## Acknowledgements

We thank Prof. Bobbi Helgason for her support and Alix Schebel for her technical assistance in sample collection, processing and DNA extraction. We also acknowledge the University of Saskatchewan for providing access to high-performance computing resources.

## Author contributions

**YL**: Writing - original draft; **SDS**, **SV**, and **YL**: Writing - review and editing; **SDM and YL**: Data curation; **YL**: Formal analysis; **YL**: Visualization; **SDM** and **SDS**: Methodology; **SDS**: Conceptualization; **SDS**: Resources; **SDS**: Funding acquisition; **SDS**: Project administration; **SDS**: Supervision.

## Conflict of interest

No conflict of interest declared

## Funding

This study was supported by the Plant Phenotyping and Imaging Research Centre (P_2_IRC) at the University of Saskatchewan, Canada.

## Data availability

Shotgun metagenomic sequencing data were deposited on the National Center for Biotechnology Information (BioProject, accession number PRJNA1104571)

## Notes

### Competing Interest Statement

The authors have declared no competing interest.

## References

Agler MT, Ruhe J, Kroll S, Morhenn C, Kim ST, Weigel D, Kemen EM. 2016. Microbial hub taxa link host and abiotic factors to plant microbiome variation. PLOS Biology 14, e1002352. doi:10.1371/journal.pbio.1002352

Barraza A, Vizuet-de-Rueda JC, Alvarez-Venegas R. 2020. Highly diverse root endophyte bacterial community is driven by growth substrate and is plant genotype-independent in common bean (Phaseolus vulgaris L.). PeerJ 8, e9423. doi:10.7717/peerj.9423

Bazghaleh N, Mamet SD, Bell JK, Moreira ZM, Taye ZM, Williams S, Arcand M, Lamb EG, Shirtliffe S, Vail S, Siciliano SD, Helgason B. 2020. An intensive multilocation temporal dataset of fungal communities in the root and rhizosphere of Brassica napus. Data in Brief 30, 105467. doi:10.1016/j.dib.2020.105467

Bergelson J, Brachi B, Roux F, Vailleau F. 2021. Assessing the potential to harness the microbiome through plant genetics. Current Opinion in Biotechnology, Food Biotechnology • Plant Biotechnology 70, 167–173. doi:10.1016/j.copbio.2021.05.007

Beschoren da Costa P, Benucci GMN, Chou MY, Van Wyk, J, Chretien M, Bonito G. 2022 Soil origin and plant genotype modulate switchgrass aboveground productivity and root microbiome assembly. mBio 0, e00079–22. doi:10.1128/mbio.00079-22

Bolger AM, Lohse M, Usadel B. 2014. Trimmomatic: a flexible trimmer for Illumina sequence data. Bioinformatics 30, 2114–2120. doi:10.1093/bioinformatics/btu170

Bonito G, Reynolds H, Robeson MS, Nelson J, Hodkinson BP, Tuskan G, Schadt CW, Vilgalys R. 2014. Plant host and soil origin influence fungal and bacterial assemblages in the roots of woody plants. Molecular Ecology 23, 3356–3370. doi:10.1111/mec.12821

Broad Institute. 2019. Picard toolkit. Broad Institute, GitHub Repository.

Buchfink B, Reuter K, Drost HG. 2021. Sensitive protein alignments at tree-of-life scale using DIAMOND. Nature Methods 18, 366–368. doi:10.1038/s41592-021-01101-x

Byers AK, Condron LM, O’Callaghan M, Waller L, Dickie IA, Wakelin SA. 2023. Plant species identity and plant-induced changes in soil physicochemistry—but not plant phylogeny or functional traits - shape the assembly of the root-associated soil microbiome. FEMS Microbiology Ecology 99, fiad126. doi:10.1093/femsec/fiad126

Coyte KZ, Rakoff-Nahoum S. 2019. Understanding competition and cooperation within the mammalian gut microbiome. Current Biology 29, R538–R544. doi:10.1016/j.cub.2019.04.017

Csardi G, Nepusz T. 2005. The Igraph software package for complex network research. InterJournal Complex Systems, 1695.

Danecek P, Bonfield JK, Liddle J, Marshall J, Ohan V, Pollard MO, Whitwham A, Keane T, McCarthy SA, Davies RM, Li H. 2021. Twelve years of SAMtools and BCFtools. GigaScience 10, giab008. doi:10.1093/gigascience/giab008

Deng, S., Caddell, D.F., Xu, G., Dahlen, L., Washington, L., Yang, J., Coleman-Derr, D., 2021. Genome wide association study reveals plant loci controlling heritability of the rhizosphere microbiome. The ISME Journal 15, 3181–3194. doi:10.1038/s41396-021-00993-z

Durán P, Thiergart T, Garrido-Oter R, Agler M, Kemen E, Schulze-Lefert P, Hacquard S. 2018. Microbial Interkingdom Interactions in Roots Promote Arabidopsis Survival. Cell 175, 973–983.e14. doi:10.1016/j.cell.2018.10.020

Ebersbach J, Khan NA, McQuillan I, Higgins EE, Horner K, Bandi V, Gutwin C, Vail LS, Robinson SJ, Parkin IA. 2022. Exploiting high-throughput indoor phenotyping to characterize the founders of a structured B. napus breeding population. Frontiers in Plant Science, 12, 780250. doi10.3389/fpls.2021.780250

Efron B. 2005. Local false discovery rates. Stanford University.

Fan W, Tang F, Wang J, Dong J, Xing J, Shi F. 2023. Drought-induced recruitment of specific root-associated bacteria enhances adaptation of alfalfa to drought stress. Frontiers in Microbiology 14.

Faust K, Raes J. 2012. Microbial interactions: from networks to models. Nature Reviews Microbiology 10, 538-550. doi:10.1038/nrmicro2832

Favela A, O Bohn M, D Kent A. 2021. Maize germplasm chronosequence shows crop breeding history impacts recruitment of the rhizosphere microbiome. The ISME Journal 15, 2454–2464. doi:10.1038/s41396-021-00923-z

Fuhrman JA. 2009. Microbial community structure and its functional implications. Nature 459, 193–199. doi:10.1038/nature08058

Hevey D. 2018. Network analysis: a brief overview and tutorial. Health Psychology and Behavioral Medicine 6, 301-328. doi:10.1080/21642850.2018.1521283

Hoek TA, Axelrod K, Biancalani T, Yurtsev EA, Liu J, Gore J. 2016. Resource availability modulates the cooperative and competitive nature of a microbial cross-feeding mutualism. PLOS Biology 14, e1002540. doi:10.1371/journal.pbio.1002540

Horton MW, Bodenhausen N, Beilsmith K, Meng D, Muegge BD, Subramanian S, Vetter MM, Vilhjálmsson BJ, Nordborg M, Gordon JI, Bergelson J. 2014. Genome-wide association study of Arabidopsis thaliana leaf microbial community. Nature Communications 5, 5320. doi:10.1038/ncomms6320

Hyatt D, Chen GL, LoCascio PF, Land ML, Larimer FW, Hauser LJ. 2010. Prodigal: prokaryotic gene recognition and translation initiation site identification. BMC Bioinformatics 11, 119. doi:10.1186/1471-2105-11-119

Ji N, Liang D, Clark LV, Sacks EJ, Kent AD. 2023. Host genetic variation drives the differentiation in the ecological role of the native Miscanthus root-associated microbiome. Microbiome 11,216. doi:10.1186/s40168-023-01646-3

Kolde R. 2019. pheatmap: Pretty Heatmap. R Package Version 1.0.12.

Kong WL, Wang WY, Zuo SH, Wu XQ. 2022. Genome sequencing of Rahnella victoriana JZ-GX1 provides new insights into molecular and genetic mechanisms of plant growth promotion. Frontiers in Microbiology 13. doi:10.3389/fmicb.2022.828990

Landa-Acuña D, Toro M, Santos-Mendoza R, Zuniga-Davila D. 2023. Role of Rahnella aquatilis AZO16M2 in phosphate solubilization and ex vitro acclimatization of Musa acuminata var. valery. Microorganisms 11, 1596. doi:10.3390/microorganisms11061596

Langmead B, Salzberg SL. 2012. Fast gapped-read alignment with Bowtie 2. Nature Methods 9, 357–359. doi:10.1038/nmeth.1923

Li H, La S, Zhang X, Gao L, Tian Y. 2021. Salt-induced recruitment of specific root-associated bacterial consortium capable of enhancing plant adaptability to salt stress. The ISME Journal 15, 2865–2882. doi:10.1038/s41396-021-00974-2

Li Y, Bazghaleh N, Vail S, Mamet SD, Siciliano SD, Helgason B. 2023a. Root and rhizosphere fungi associated with the yield of diverse Brassica napus genotypes. Rhizosphere 25, 100677. doi:10.1016/j.rhisph.2023.100677

Li Y, Vail SL, Arcand MM, Helgason BL. 2023b. Contrasting nitrogen fertilization and Brassica napus (Canola) variety development impact recruitment of the root-associated microbiome. Phytobiomes Journal 7, 125–137. doi:10.1094/PBIOMES-07-22-0045-R

Lin H, Peddada SD. 2024. Multigroup analysis of compositions of microbiomes with covariate adjustments and repeated measures. Nature Methods 21, 83–91. doi:10.1038/s41592-023-02092-7

Lin H, Peddada SD. 2020a. Analysis of compositions of microbiomes with bias correction. Nature Communications 11,3514. doi:10.1038/s41467-020-17041-7

Lin H, Peddada SD. 2020b. Analysis of microbial compositions: a review of normalization and differential abundance analysis. Npj Biofilms and Microbiomes 6, 1–13. doi:10.1038/s41522-020-00160-w

Lo CC, Chain PSG. 2014. Rapid evaluation and quality control of next generation sequencing data with FaQCs. BMC Bioinformatics 15, 366. doi:10.1186/s12859-014-0366-2

Lu J, Breitwieser FP, Thielen P, Salzberg SL. 2017. Bracken: estimating species abundance in metagenomics data. PeerJ Computer Science 3, e104. doi:10.7717/peerj-cs.104

Lu J, Rincon N, Wood DE, Breitwieser FP, Pockrandt C, Langmead B, Salzberg SL, Steinegger M. 2022. Metagenome analysis using the Kraken software suite. Nature Protocols 17, 2815–2839. doi:10.1038/s41596-022-00738-y

Mamet SD, Helgason BL, Lamb EG, McGillivray A, Stanley KG, Robinson SJ, Aziz SU, Vail S, Siciliano SD. 2021. Phenology-dependent root bacteria enhance yield of Brassica napus. Soil Biology and Biochemistry 108468. doi:10.1016/j.soilbio.2021.108468

Meng L, Xu C, Wu F. 2022. Microbial co-occurrence networks driven by low-abundance microbial taxa during composting dominate lignocellulose degradation. Science of The Total Environment 845, 157197. doi:10.1016/j.scitotenv.2022.157197

Morales Moreira ZP, Helgason BL, Germida JJ. 2021. Environment has a stronger effect than host plant genotype in shaping spring Brassica napus seed microbiomes. Phytobiomes Journal 5, 220–230. doi:10.1094/PBIOMES-08-20-0059-R

Myszka K, Tomas N, Wolko Ł. 2023. Gallic and ferulic acids suppress proteolytic activities and volatile trimethylamine production in the food-borne spoiler Rahnella aquatilis KM05. Journal of the Science of Food and Agriculture 103, 6584–6594. doi:10.1002/jsfa.12753

Nurk S, Meleshko D, Korobeynikov A, Pevzner PA. 2017. metaSPAdes: a new versatile metagenomic assembler. Genome Research 27, 824–834. doi:10.1101/gr.213959.116

Ofek M, Voronov-Goldman M, Hadar Y, Minz D. 2014. Host signature effect on plant root-associated microbiomes revealed through analyses of resident vs. active communities. Environmental Microbiology 16, 2157–2167. doi:10.1111/1462-2920.12228

Pang Z, Chen J, Wang T, Gao C, Li Z, Guo L, Xu J, Cheng Y. 2021. Linking plant secondary metabolites and plant microbiomes: A Review. Frontiers in Plant Science 12.

Patro R, Duggal G, Love MI, Irizarry RA, Kingsford C. 2017. Salmon provides fast and bias-aware quantification of transcript expression. Nature Methods 14, 417–419. doi:10.1038/nmeth.4197

Peng J, Wu D, Liang Y, Li L, Guo Y. 2019. Disruption of acdS gene reduces plant growth promotion activity and maize saline stress resistance by Rahnella aquatilis HX2. Journal of Basic Microbiology 59, 402–411. doi:10.1002/jobm.201800510

Pérez-Jaramillo JE, Carrión VJ, Bosse M, Ferrão LFV, de Hollander M, Garcia AAF, Ramírez CA, Mendes R, Raaijmakers JM. 2017. Linking rhizosphere microbiome composition of wild and domesticated Phaseolus vulgaris to genotypic and root phenotypic traits. The ISME Journal 11, 2244–2257. doi:10.1038/ismej.2017.85

Peschel S, Müller CL, von Mutius E, Boulesteix AL, Depner M. 2021. NetCoMi: network construction and comparison for microbiome data in R. Briefings in Bioinformatics 22, bbaa290. doi:10.1093/bib/bbaa290

Quinlan AR, Hall IM. 2010. BEDTools: a flexible suite of utilities for comparing genomic features. Bioinformatics 26, 841–842. doi:10.1093/bioinformatics/btq033

Reeves TG, Cassaday K. 2002. History and past achievements of plant breeding. Australian Journal of Agricultural Research 53, 851–863. doi:10.1071/ar02038

Ruhnau B. 2000. Eigenvector-centrality — a node-centrality? Social Networks 22, 357–365. doi:10.1016/S0378-8733(00)00031-9

Schmitz L, Yan Z, Schneijderberg M, de Roij M, Pijnenburg R, Zheng Q, Franken C, Dechesne A, Trindade LM, van Velzen R, Bisseling T, Geurts R, Cheng X. 2022. Synthetic bacterial community derived from a desert rhizosphere confers salt stress resilience to tomato in the presence of a soil microbiome. The ISME Journal 16, 1907–1920. doi:10.1038/s41396-022-01238-3

Segata N, Izard J, Waldron L, Gevers D, Miropolsky L, Garrett WS, Huttenhower C. 2011. Metagenomic biomarker discovery and explanation. Genome Biology 12, R60. doi:10.1186/gb-2011-12-6-r60

Simonin M, Dasilva C, Terzi V, Ngonkeu ELM, Diouf D, Kane A, Béna G, Moulin L. 2020. Influence of plant genotype and soil on the wheat rhizosphere microbiome: evidences for a core microbiome across eight African and European soils. FEMS Microbiology Ecology 96, fiaa067. doi:10.1093/femsec/fiaa067

Singh KD, Duddu HSN, Vail S, Parkin I, Shirtliffe SJ. 2021. UAV-based hyperspectral imaging technique to estimate canola (Brassica napus L.) seedpods maturity. Canadian Journal of Remote Sensing 47, 33–47. doi:10.1080/07038992.2021.1881464

Su F, Zhao B, Dhondt-Cordelier S, Vaillant-Gaveau N. 2024. Plant-growth-promoting rhizobacteria modulate carbohydrate metabolism in connection with host plant defense mechanism. International Journal of Molecular Sciences 25, 1465. doi:10.3390/ijms25031465

Tapio I, Fischer D, Blasco L, Tapio M, Wallace RJ, Bayat AR, Ventto L, Kahala M, Negussie E, Shingfield KJ, Vilkki J. 2017. Taxon abundance, diversity, co-occurrence and network analysis of the ruminal microbiota in response to dietary changes in dairy cows. PLOS ONE 12, e0180260. doi:10.1371/journal.pone.0180260

Taye ZM, Helgason BL, Bell JK, Norris CE, Vail S, Robinson SJ, Parkin IAP, Arcand M, Mamet S, Links MG, Dowhy T, Siciliano S, Lamb EG. 2020. Core and differentially abundant bacterial taxa in the rhizosphere of field grown Brassica napus genotypes: implications for canola breeding. Frontiers in Microbiology 10, 3007. doi:10.3389/fmicb.2019.03007

Tenenbaum D, Bioconductor Package Maintainer. 2023. KEGGREST: Client-side REST access to the Kyoto Encyclopedia of Genes and Genomes (KEGG). R Package Version 1.40.1. doi:10.18129/B9.bioc.KEGGREST

Tu Q, Lin L, Cheng L, Deng Y, He Z. 2019. NCycDB: a curated integrative database for fast and accurate metagenomic profiling of nitrogen cycling genes. Bioinformatics 35, 1040–1048. doi:10.1093/bioinformatics/bty741

Vollmers J, Wiegand S, Lenk F, Kaster AK. 2022. How clear is our current view on microbial dark matter? (Re-)assessing public MAG & SAG datasets with MDMcleaner. Nucleic Acids Research 50, e76. doi:10.1093/nar/gkac294

Wagner MR, Lundberg DS, del Rio TG, Tringe SG, Dangl JL, Mitchell-Olds T. 2016. Host genotype and age shape the leaf and root microbiomes of a wild perennial plant. Nature Communications 7, 12151. doi:10.1038/ncomms12151

Wang B, Jin H, Xu Y, Sun Z. 2023. Isolation, characterization, and genomic analysis of multidrug-resistant Rahnella aquatilis from fruits in China. Current Microbiology 80, 321. doi:10.1007/s00284-023-03436-4

Wang C, Li Y, Li M, Zhang K, Ma W, Zheng L, Xu H, Cui B, Liu R, Yang Y, Zhong Y, Liao H. 2021. Functional assembly of root-associated microbial consortia improves nutrient efficiency and yield in soybean. Journal of Integrative Plant Biology 63, 1021–1035. doi:10.1111/jipb.13073

Williams ST. 2023. Linking Soil Nitrogen cycling and plant biotic traits to nitrogen use efficiency parameters among diverse canola (Brassica napus) genotypes. [Doctoral dissertation, University of Saskatchewan] Https://harvest.usask.ca/bitstreams/f9e1210d-d88a-4217-bac0-413c130e498f/download

Williams ST, Vail S, Arcand MM. 2021. Nitrogen use efficiency in parent vs. hybrid canola under varying nitrogen availabilities. Plants 10, 2364. doi:10.3390/plants10112364

Wood DE, Lu J, Langmead B. 2019. Improved metagenomic analysis with Kraken 2. Genome Biology 20, 257. doi:10.1186/s13059-019-1891-0

Wood DE, Salzberg SL. 2014. Kraken: ultrafast metagenomic sequence classification using exact alignments. Genome Biology 15, R46. doi:10.1186/gb-2014-15-3-r46

Xu S, Zhao Y, Peng Y, Shi Y, Xie X, Chai A, Li B, Li L. 2022. Comparative genomics assisted functional characterization of Rahnella aceris ZF458 as a novel plant growth promoting Rhizobacterium. Frontiers in Microbiology 13. doi:10.3389/fmicb.2022.850084

Yin C, Casa Vargas JM, Schlatter DC, Hagerty CH, Hulbert SH, Paulitz TC. 2021. Rhizosphere community selection reveals bacteria associated with reduced root disease. Microbiome 9, 86. doi:10.1186/s40168-020-00997-5

Yu, P., He, X., Baer, M., Beirinckx, S., Tian, T., Moya, Y.A.T., Zhang, X., Deichmann, M., Frey, F.P., Bresgen, V., Li, C., Razavi BS, Schaaf G, von Wirén N, Su Z, Bucher M, Tsuda K, Goormachtig S, Chen X, Hochholdinger F. 2021. Plant flavones enrich rhizosphere Oxalobacteraceae to improve maize performance under nitrogen deprivation. Nature Plants 7, 481–499. doi:10.1038/s41477-021-00897-y

Zeng J, Tu Q, Yu X, Qian L, Wang C, Shu L, Liu F, Liu S, Huang Z, He J, Yan Q, He Z. 2022. PCycDB: a comprehensive and accurate database for fast analysis of phosphorus cycling genes. Microbiome 10, 101. doi:10.1186/s40168-022-01292-1

Zhang J, Liu YX, Zhang N, Hu B, Jin T, Xu H, Qin Y, Yan P, Zhang X, Guo X, Hui J, Cao S, Wang X, Wang C, Wang H, Qu B, Fan G, Yuan L, Garrido-Oter R, Chu C, Bai Y. 2019. NRT1.1B is associated with root microbiota composition and nitrogen use in field-grown rice. Nature Biotechnology 37, 676–684. doi:10.1038/s41587-019-0104-4

Zhang L, Feng G, Declerck S. 2018. Signal beyond nutrient, fructose, exuded by an arbuscular mycorrhizal fungus triggers phytate mineralization by a phosphate solubilizing bacterium. The ISME Journal 12, 2339–2351. doi:10.1038/s41396-018-0171-4

Zhang T, Vail S, Duddu HSN, Parkin IAP, Guo X, Johnson EN, Shirtliffe SJ. 2021. Phenotyping flowering in canola (Brassica napus L.) and estimating seed yield using an unmanned aerial vehicle-based imagery. Frontiers in Plant Science 12.

Zheng J, Ge Q, Yan Y, Zhang X, Huang L, Yin Y. 2023. dbCAN3: automated carbohydrate-active enzyme and substrate annotation. Nucleic Acids Research 51, W115–W121. doi:10.1093/nar/gkad328

Zheng J, Huang L, Yi H, Yan Y, Zhang X, Akresi J, Yin Y. 2024. Carbohydrate-active enzyme annotation in microbiomes using dbCAN. doi:10.1101/2024.01.10.575125

Zhou H, He K, Chen J, Zhang X. 2022. LinDA: linear models for differential abundance analysis of microbiome compositional data. Genome Biology 23, 95. doi:10.1186/s13059-022-02655-5

Zouagui R, Zouagui H, Aurag J, Ibrahimi A, Sbabou L. 2024. Functional analysis and comparative genomics of Rahnella perminowiae S11P1 and Variovorax sp. S12S4, two plant growth-promoting rhizobacteria isolated from Crocus sativus L. (saffron) rhizosphere. BMC Genomics 25, 289. doi:10.1186/s12864-024-10088-6

